# Heterologous expression of the cyanobacterial fructose-1,6-/sedoheptulose-1,7-bisphosphatase in *Chlamydomonas reinhardtii* causes increased cell size and biomass productivity in mixotrophic conditions

**DOI:** 10.1101/2025.05.02.651950

**Authors:** Martina Bussola, Federico Perozeni, Maria Meloni, Matteo Pivato, Mirko Zaffagnini, Matteo Ballottari

## Abstract

The Calvin-Benson-Bassham (CBB) cycle is the metabolic pathway responsible for CO assimilation in oxygenic photosynthetic organisms. Two key rate-limiting steps in this cycle are catalyzed by the enzymes fructose-1,6-bisphosphatase (FBPase) and sedoheptulose-1,7-bisphosphatase (SBPase), making them promising targets for genetic enhancement to improve carbon fixation. A potential strategy involves overexpressing a cyanobacterial dual-function FBP/SBPase, which catalyzes both reactionsOverexpression of this enzyme in tobacco plants or in other plants led to an increase in growth rate and biomass accumulation. Here, the overexpression of the same enzyme was achieved in *Chlamydomonas reinhardtii*. The recombinant cyanobacterial FBP/SBPase isolated from *C. reinhardtii* exhibited the expected catalytic activity, being Mg^2+^ dependent and strongly activated in the presence of a reducing agent. The FBP/SBPase expressing lines exhibited an increased photosynthetic activity at the cell level and decreased production of singlet oxygen upon exposure to high irradiances, suggesting improved capacity to manage high excitation pressure of the photosynthetic apparatus. Increased cell volume was measured in FBP/SBPase-expressing lines under different growth conditions. However, increased growth and biomass productivity were observed only in mixotrophy when light and CO2 were limiting, leading to increased starch, protein, and lipid content on a cellular basis. This phenotype caused an increased sedimentation rate in the transformant lines: the expression of FBP/SBPase enzyme could thus be considered as a strategy to improve the cell harvesting process. These findings provide new insights into carbon metabolism in microalgae, and could, in the future, support improved biomass accumulation, paving the way for effective domestication and industrial use.

## 1. INTRODUCTION

Microalgae are photosynthetic organisms with a broad range of potential applications for their biomass and derivatives, spanning from food and feed to the production of high-value products of pharmaceutical and nutraceutical interest ^1–8^. From an industrial point of view, biomass production needs to be increased to reduce cultivation cost and improve yield ^9^. The purpose of obtaining algae strains with higher productivity can be achieved by genetic manipulation of various metabolic pathways, including carbon fixation ^10,11^. In eukaryotic photosynthetic organisms, the metabolic phase of photosynthesis takes place in the chloroplast stroma, where NADPH and ATP, synthesized in the light phase, are used to drive carbon assimilation through the Calvin-Benson-Bassham (CBB) cycle. This biochemical pathway proceeds with three different and coordinated phases: carboxylation, reduction, and regeneration. In the carboxylation phase, Rubisco catalyzes the fixation of inorganic carbon (*i.e.*, CO_2_) onto a pentose acceptor (ribulose-1,5-bisphosphate, RuBP), generating two molecules of a three-carbon intermediate (3-phosphoglyceric acid, 3-PGA). In the reduction phase, 3-PGA is sequentially converted to glyceraldehyde-3-phosphate (G3P) through the action of phosphoglycerate kinase (PGK) and glyceraldehyde-3-phosphate dehydrogenase (GAPDH). This process consumes ATP and NADPH, serving as sources of energy and reducing power, respectively. The cycle is completed in the third phase by the regeneration of RuBP, the substrate of Rubisco: this phase involved eight enzymes and requires ATP as the sole energy source ^12^. The coordination between the photosynthetic light reactions and the CBB cycle is governed by light-dependent redox-regulatory mechanisms. These mechanisms operate primarily through the ferredoxin/thioredoxin system, which control the activities of several CBB cycle enzymes via reversible disulfide/dithiol exchange reactions, thereby enabling dynamic regulation of carbon fixation in response to the cellular redox state ^13–15^.

Given its central role in carbon assimilation and its direct impact on biomass production, CBB cycle represents a highly promising target for metabolic engineering to improve carbon fixation. Multiple studies have investigated the factors limiting the overall efficiency of the CBB cycle, consistently identifying Rubisco as the most significant bottleneck ^11,12,16–20^. This is primarily due to its relatively slow catalytic rate and dual carboxylase/oxygenase activity, which can lead to energetically wasteful photorespiration ^21,22^. In addition to Rubisco, two enzymes involved in the regeneration phase of the CBB cycle, sedoheptulose-1,7-bisphosphatase (SBPase) and fructose-1,6-bisphosphatase (FBPase), have also been identified as limiting steps in this metabolic pathway ^11,12,20,23–25^. The constraints associated with FBPase- and SBPase-catalyzed reactions have been addressed by modulating the expression levels of these enzymes and evaluating the impact on the overall efficiency of the CBB cycle ^10,11,23,25^. FBPase is an enzyme broadly distributed across animals, plants, and microorganisms. In contrast, SBPase plays a unique role in the CBB cycle and is exclusively found in photoautotrophic organisms. At the kinetic level, both enzymes catalyze irreversible dephosphorylation reactions. FBPase specifically hydrolyzes fructose-1,6-bisphosphate (FBP) to generate fructose-6-phosphate (F6P) and inorganic phosphate, whereas SBPase acts on sedoheptulose-1,7-bisphosphate (SBP), producing sedoheptulose-7-phosphate (S7P) and inorganic phosphate ^12^. Chloroplast FBPase and SBPase functions have been studied for years in plants such as wheat, potato, tobacco, Arabidopsis, pea and many others: according to these studies, light is a key regulator of these enzymes for both gene expression and protein catalytic activity ^25–31^. *In vitro* studies have further confirmed that FBPase and SBPase are redox-regulated enzymes, exhibiting higher activity under reducing conditions compared to oxidizing conditions ^32–37^. This regulation is attributed to the formation/reduction of disulfide bonds between conserved cysteine residues that modulate their activity ^32,33,36^.

In the model green microalga *Chlamydomonas reinhardtii*, FBPase is encoded by two nuclear genes, FBP1 and FBP2, which give rise to two isoforms localized in distinct subcellular compartments ^38,39^. The cytoplasmic FBPase functions primarily in gluconeogenesis, while the chloroplast isoform participates in the regeneration of RuBP within the CBB cycle and in the starch synthesis pathway ^39^. In contrast, SBPase is encoded by a single nuclear gene in *Chlamydomonas reinhardtii* (SBP1) and is localized exclusively in the chloroplast ^36^.

*C. reinhardtii* has been exploited as biological chassis to test the impact of overexpressing FBPase and SBPase as a possible strategy to improve carbon fixation ^10,38^. Dejtisakdi and colleagues described the transformation of *C. reinhardtii* chloroplast to overexpress the cyanobacterial FBPase and observed a 1.4-fold increase in total FBPase activity ^38^. However, the growth of the mutant line was very similar to that of parental strain CC125 under mixotrophic conditions and atmospheric CO_2_, while under photoautotrophic growth conditions or with elevated CO_2_ levels, a negative effect on growth rate and biomass production was observed ^38^. Although this result is in contrast with what was previously observed in higher plants, FBPase probably does not catalyze a rate limiting step in *C. reinhardtii*, likely due to the highly complex regulation of the CBB cycle and its central role in cellular metabolism. The overexpression of SBPase in *C. reinhardtii* was also reported demonstrating that increased expression of endogenous SBPase enhances photosynthetic performance and promotes biomass accumulation in cells exposed to 150 µmol photons m^−2^ s^−1^ and elevated CO_2_ concentrations ^10^. These findings are consistent with previous observations, suggesting that SBPase abundance constitutes a key bottleneck in the metabolic flux through the CBB cycle in both microalgae and terrestrial plants ^12,24,25,27,40^.

In certain cyanobacterial species, such as *Synechococcus* PCC 7942, FBPase enzymes exhibiting dual FBPase/SBPase activity (*i.e.*, dephosphorylation of FBP and SBP) have been identified^41^. The expression of this dual-function cyanobacterial enzyme has been proposed as a potential strategy to enhance carbon fixation efficiency: Miyagawa and colleagues reported that the overexpression of a cyanobacterial FBP/SBPase in the chloroplasts of transgenic *Nicotiana tabacum cv. Xanth* plants resulted in enhanced photosynthetic capacity, increased carbohydrate accumulation, and accelerated growth rate ^28^. In particular, they demonstrated that, under atmospheric conditions (360 p.p.m. CO_2_), final dry matter and photosynthetic CO_2_ fixation of the transgenic plants were 1.5-fold and 1.24-fold higher, respectively^28^. It was also reported that transgenic tobacco plants exhibited a 1.2-fold increase in the initial activity of Rubisco compared to wild-type plants. Additionally, levels of CBB cycle intermediates were significantly higher in the transgenic lines ^28^.

Additional studies in plant species such as lettuce, soybean, and rice, as well as in the microalga *Euglena gracilis*, have supported the positive impact of this dual-function enzyme on enhancing photosynthetic rate and promoting growth ^42–45^. However, it is important to note that the balance between different steps of the CBB cycle can vary significantly among species: as mentioned, only the overexpression of SBPase enzyme in *C. reinhardtii* led to improved biomass productivity ^10^, while FBPase overexpression had a limited or even negative effect on biomass productivity ^38^.

This work aims to express the cyanobacterial FBPase/SBPase gene in *Chlamydomonas reinhardtii* to tune carbon fixation metabolism. To our knowledge, the expression of this cyanobacterial dual FBP/SBPase in *C. reinhardtii* has not yet been reported in the literature. Overexpression of FBP/SBPase in *C. reinhardtii* increased cell size and cellular photosynthetic activity, either in photoautotrophic or mixotrophic conditions. Moreover, the expression of cyanobacterial FBP/SBPase resulted in faster growth rate in mixotrophy under limiting light and low CO_2_ availability.

## 2. MATERIALS AND METHODS

### 2.1 Cloning Procedure, Transformation, and Colony Screening

The strain *Chlamydomonas reinhardtii* UVM4 ^46^ was used as the background strain for nuclear engineering. Codon-optimized coding sequences (CDS) for FBP/SBPase from *Synechococcus elongatus* (Q31QY2) was inserted into the pOpt2 expression vector ^47–49^. Additionally, sequences from the RBCS2 intron 1 were included to enhance expression in *Chlamydomonas* as described previously ^47,50^. PsaD (Photosystem I subunit D) transit peptide was used to drive protein localization in the chloroplast. Hybrid HSP70/RBCS promoter was used for gene expression, being previously described as a strong promoter for *C. reinhardtii* ^51^. Two different constructs were generated, respectively, with the presence (Psad_FBP/SBPase_YFP) or absence (Psad_FBP/SBPase) of YFP coding sequence (mVenus) downstream of the FBP/SBPase CDS to generate a fusion protein. The mVenus sequence in the expression vector contained the RBCS2 intron 2 sequence to maximize gene expression as previously reported ^47,50^. Fusion of the FBP/SBPase to the fluorescent protein was used to enable protein localization. Cloning was carried out using Thermo FastDigest restriction enzymes, followed by ligation into pOpt2 vectors. s. Transformation into *C. reinhardtii* was performed with the glass beads method using 10 µg of vector DNA linearized with XbaI and KpnI restriction enzymes ^47^. Transformants were selected by plating on TAP agar containing 12 mg/L paromomycin. In the case of transformation with Psad_FBP/SBPase_YFP construct antibiotic-resistant colonies were first screened for fluorescence (excitation 509 ± 9 nm, emission 540 ± 20 nm), and the brightest fluorescent lines were isolated ^47^. These lines were cultured individually and subjected to SDS-PAGE analysis, followed by western blotting and immunodetection using an antibody recognizing the YFP protein. In the case of transformation with Psad_FBP/SBPase construct antibiotic-resistant colonies were directly screened by western blot and immunodetection using an antibody recognizing Strep-tag peptide placed at the C-terminus of FBO/SBPase protein. In both cases, positive colonies, identified by a band at the expected molecular weight (38 and 65 kDa for PsaD_FBP/SBPase and PsaD_FBP/SBPase_YFP respectively), were selected for further analysis. Isolated expressing lines were maintained on TAP agar plates under continuous white LED light set at 40 μmol photons m^-2^ s^-1^.

### 2.2 Fluorescence microscopy and subcellular localization

Subcellular localization of PsaD_FBP/SBPase_YFP was assessed via confocal microscopy as described in previous studies. Imaging was conducted using a Leica TCS-SP5 inverted confocal microscope (Leica Microsystems, Germany). For fluorescence detection of YFP, the excitation wavelength was 524 nm and Emission from 522 to 572 nm was recorded. For chlorophyll fluorescence, the excitation wavelength was 524 nm, with detection occurring in the 680-720 nm range.

### 2.3 Microalgae cultivation

*Chlamydomonas reinhardtii* UVM4 and engineered lines were grown in High-Salt (HS) or Tris-acetate-phosphate (TAP) media ^52,53^ . Growth experiments were performed in flasks exposed to 100 µmol photons m^−2^ s^−1^ or in small scale airlifted photobioreactor with a volume of 80 mL in the Multi-Cultivator MC 1000-OD (Photon System Instrument) system. The growth experiments in Multi-Cultivator MC 1000-OD were performed in tubes aerated with air or with 3% CO_2_-enriched air obtained by a gas mixing system (Gas Mixing System GMS 150, Photon System Instrument). In the Multi-Cultivator MC 1000-OD cells were exposed to different irradiances (100 or 1000 µmol photons m^−2^ s^−1^). Cell density and dimension were determined by Multizer 4e Coulter (Beckman Coulter).

### 2.4 Biomass analysis

Dry weight and biomass composition were analyzed at the end of the growth curves. Cells were harvested by centrifugation at 4500 x *g* for 5 minutes at 20°C and then dried in a lyophilizer for 48 h followed by gravimetric evaluation. Proteins content was analyzed by bicinchoninic acid (BCA) protein assay (Thermo Fisher Scientific). Starch content was analyzed by colorimetric reaction with iodine, as previously described^54^. Lipid content was evaluated using the fluorescent dye BODIPY 505/5015 as previously reported ^55^.

Pigments were extracted from intact cells using 80% acetone and analyzed by fitting of absorption spectra as previously described ^56^. Absorption spectra were collected using Jasco V-550 UV/VIS spectrophotometer.

### 2.5 Photosynthetic activity

The light-dependent oxygen evolution activity of the cultures was measured on samples with a cell density of 3x10^6^ cells mL^−1^ at 25 °C with a Clark-type O2 electrode (Oxygraph Plus, Hansatech) during illumination with light from a halogen lamp (Schott) at different actinic lights (from 50 to 2500 µmol photons m^−2^ s^−1^) in presence of 5 mM sodium bicarbonate ^57^. Net oxygen evolution data were obtained upon subtraction of oxygen consumption rate in the dark and fitted with the hyperbolic function y= Pmax * x/(KI + x), where Pmax is the maximum net oxygen evolution rate and KI the light intensity at which the net oxygen evolution rate is Pmax/2. Photosynthetic parameters ΦPSII, qL, electron transport rate (ETR), and NPQ^58,59^ were characterized by measuring with a DUAL-PAM-100 fluorimeter (Heinz–Walz) chlorophyll fluorescence of intact cells, at room temperature in a 1 × 1 cm cuvette mixed by magnetic stirring ^60^.

Singlet oxygen production was measured using the Singlet Oxygen Sensor Green (SOSG), a fluorescent probe that increases the intensity of its emission in the presence of this ROS ^61,62^ .Cell suspension, incubated with SOSG, were illuminated with red light at 2000 μmol photons m^−2^ s^−1^ at 20 °C. Singlet oxygen production was measured as increased fluorescence emission of SOSG compared to the initial value (excitation 480 nm, emission 510-540 nm).

### 2.6 SDS-PAGE and immunoblotting

SDS-PAGE analysis was performed using the Tris-Tricine buffer system ^63^. Immunoblotting analysis were performed using the following antibodies obtained from Agrisera company (https://www.agrisera.com/, Sweden): α-PsaA (AS06 172, dilution used 1:3000), α-PsbC (CP43) (AS11 1787, dilution used 1:2000), α-AtpC (AS08 312, dilution used 1:10000), α-RbcL (AS03 037, dilution used 1:5000), α-Lhcbm5 (AS09 408, dilution used 1:5000), α-CAH3 (AS05 073, dilution used 1:2000). For the identification of expressing lines the following antibody, obtaiend from Thermo Scientific were used: α-Strep-Tag (PA5-114454) and α-GFP (A-11122) which can recognize also the YFP. The secondary antibody used was an anti-rabbit IGG (A3687, Merck) conjugated with alkaline phosphatase as chromogenic detection system.

### 2.7 Recombinant FBP/SBPase protein extraction and quantification

Cyanobacterial FBP/SBPase recombinant protein was isolated according to the following protocol. Cells at exponential phase were harvested by centrifugation at 5000 g for 10 minutes and resuspended in 100 mM tris pH 8, 150 mM NaCl, 1 mM EDTA. Cells were then sonicated with 6 cycles of 30 seconds and loaded in an avidin-based affinity chromatography. Strep-Tactin XT Superflow High-Capacity resin column was used for protein purification. Recombinant proteins were then eluted with 100 mM Tris-HCl, pH 8, 150 mM NaCl, 1 mM EDTA, and 50 mM biotin. Quantification of the partially purified enzyme was carried out by band analysis of SDS[PAGE gel using ImageJ software. Recombinant FBPase from *Chlamydomonas reinhardtii* was used as a standard to linearly correlate protein load (1-3 μg) and band intensity. Optimal image processing was performed by background subtraction.

### 2.8 Enzymatic FBPase activity analysis

The activity of dephosphorylation of FBP was assayed spectrophotometrically by measuring the reduction of NADP^+^ at 340 nm using a Jasco V-550 UV/VIS spectrophotometer. The reaction mix (1 mL) contained 100 mM Tris-HCl, pH 7.9, 1.2 mM FBP, 0-32 mM MgSO_4_, 0.2 mM NADP^+^, 0.5 units of phosphoglucoisomerase, and 0.5 units of glucose 6-phosphate dehydrogenase ^64^. Maximal activity was obtained by incubating the purified enzyme with 20 mM dithiothreitol (DTT) for 30 min and specific activity values were normalized against protein amount. Michaelis-Menten constant of the fully active enzyme was determined by measuring the activity with variable FBP concentrations (0.05-1.2 mM) and plotted by nonlinear regression using the Michaelis-Menten equation with the GraphPad PRISM 10 software.

### 2.9 Statistics analysis

All the experiments described were performed on at least two independent lines for each genotype in at least three independent biological experiments. Statistical analysis was performed by using a two-sided Student’s t-test or one-way ANOVA with post-hoc Tukey test in the case of multiple comparisons.

## 3. RESULTS

### 3.1 Heterologous expression of cyanobacterial dual-function enzyme FBP/SBPase in Chlamydomonas reinhardtii

Overexpression of heterologous FBPase/SBPase gene in *C. reinhardtii* was achieved adopting the strategy previously developed for efficient nuclear gene expression in this species ^47,49,65^. The cyanobacterial FBP/SBPase enzyme is encoded by a gene of 1068 bp open reading frame, resulting in a polypeptide of 356 amino acid residues ^41^. In brief, the gene sequence was synthetically designed considering the codon usage of *C. reinhardtii* and inserting to the coding sequence the intron 1 of RUBISCO SMALL SUBUNIT 2 gene from *C. reinhardtii* at a specific frequency previously reported to maximize gene expression ^50,66^. PsaD target peptide was used for chloroplast localization and the strong hybrid promoter Hsp70/RBCS2 was used to drive gene expression **(Figure 1a**). The expression cassette also contained the resistance to paromomycin as a selection marker. A second expression construct was also designed, fusing at the C-terminus of FBP/SBPase enzyme the sequence encoding mVenus (YFP), creating a single fusion protein allowing visualization of protein localization by confocal microscopy ^50,66^. Colonies were selected on TAP agar plates containing paromomycin as a selection agent. In the case of resistant lines transformed with PsaD_FBPase/SBPase construct, selection of expressing lines was carried out by western blot using a primary mouse antibody that recognizes the Strep-Tag II peptide placed at the C-terminus of the protein. Four lines were found to express the protein of interest **(Figure 1b)**, representing 12.5% of the total resistant lines. In the case of lines transformed with PsaD_FBP/SBPase_YFP construct, YFP fluorescence was measured to screen the transformant lines since the fluorophore fused to the FBP/SBPase protein provides a linear correlation between protein accumulation and fluorescence signal (**Figure 1c**). The five colonies showing higher fluorescence were then analyzed by western blot using an α-GFP antibody to detect the presence of the heterologous protein.

**Figure 1.**
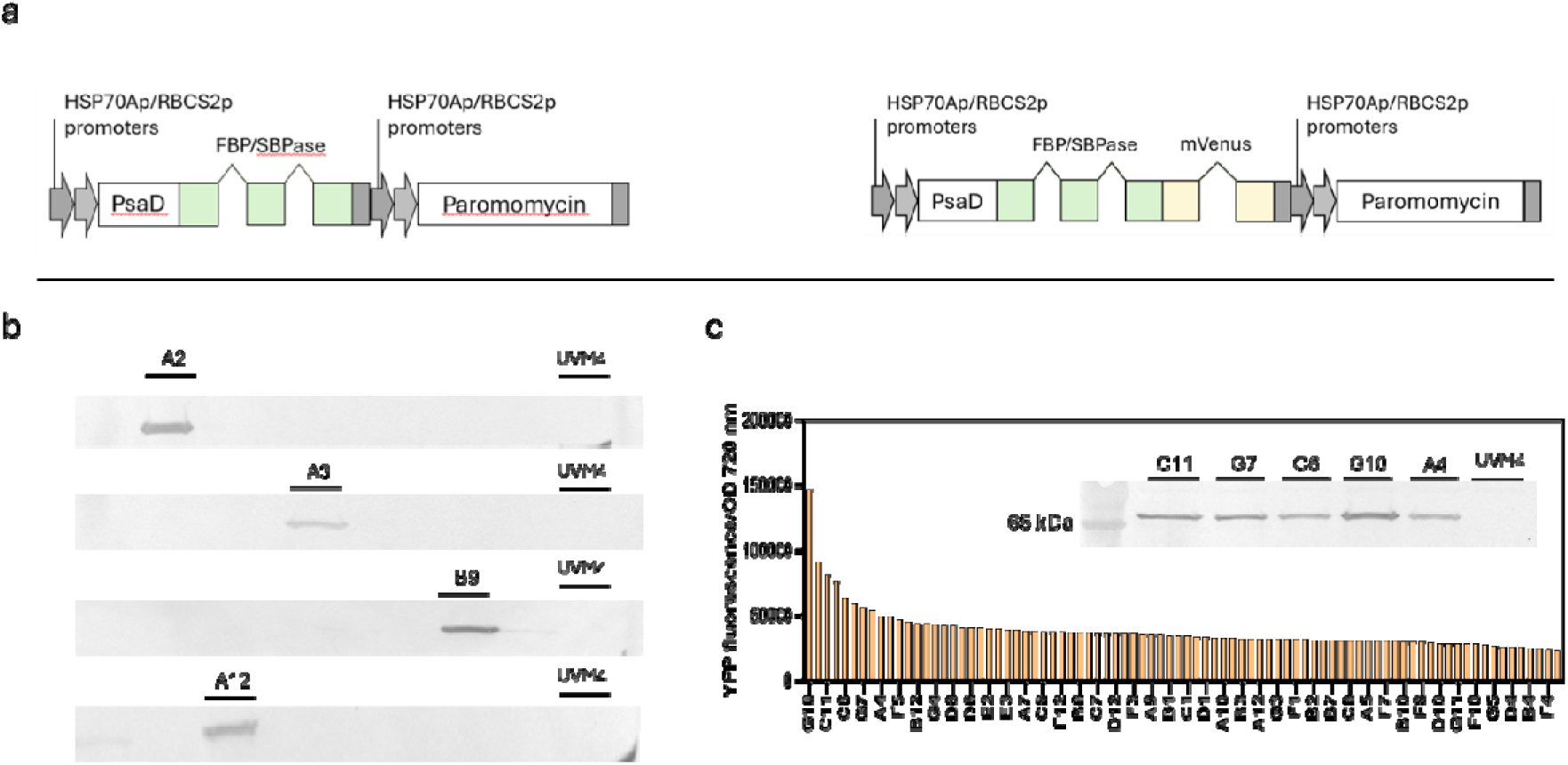
Genetic constructs and expression analysis of cyanobacterial FBP/SBPase in *C. reinhardtii.* (a) Schematic representation of expression cassettes used to generate FBP/SBPase expressing lines both with or without mVenus (YFP) fused at the C-terminus. To drive gene expression the hybrid promoter HSP70-RBCS2 wase used, while the FBP/SBPase was targeted into chloroplast by exploiting the PsaD target peptide sequence (Photosystem I Subunit D). Both expression cassettes contained the Aph VIII gene confer resistance to the antibiotic paromomycin. Both FBP/SBPase proteins either fused or not to YFP carried a C-terminal S-tag for immunodetection and for protein purification. (b) Immunoblot against S-tag showing the FBP/SBPase accumulation in transformed lines without YFP as reporter protein. (c) Fluorescence screening of transformed lines with FBP/SBPase_YFP. The best five lines were subjected to immunoblot against YFP confirming the presence of the protein are reported in the Panel c insert.

Cellular localization of recombinant FBP/SBPase enzyme was evaluated by confocal microscopy using lines transformed with the PsaD_FBPase/SBPase_YFP construct. As reported in **Figure 2**, YFP fluorescence (green) largely overlapped with chlorophyll autofluorescence (red), indicating that the recombinant FBP/SBPase enzyme is localized in the chloroplast.

**Figure 2.**
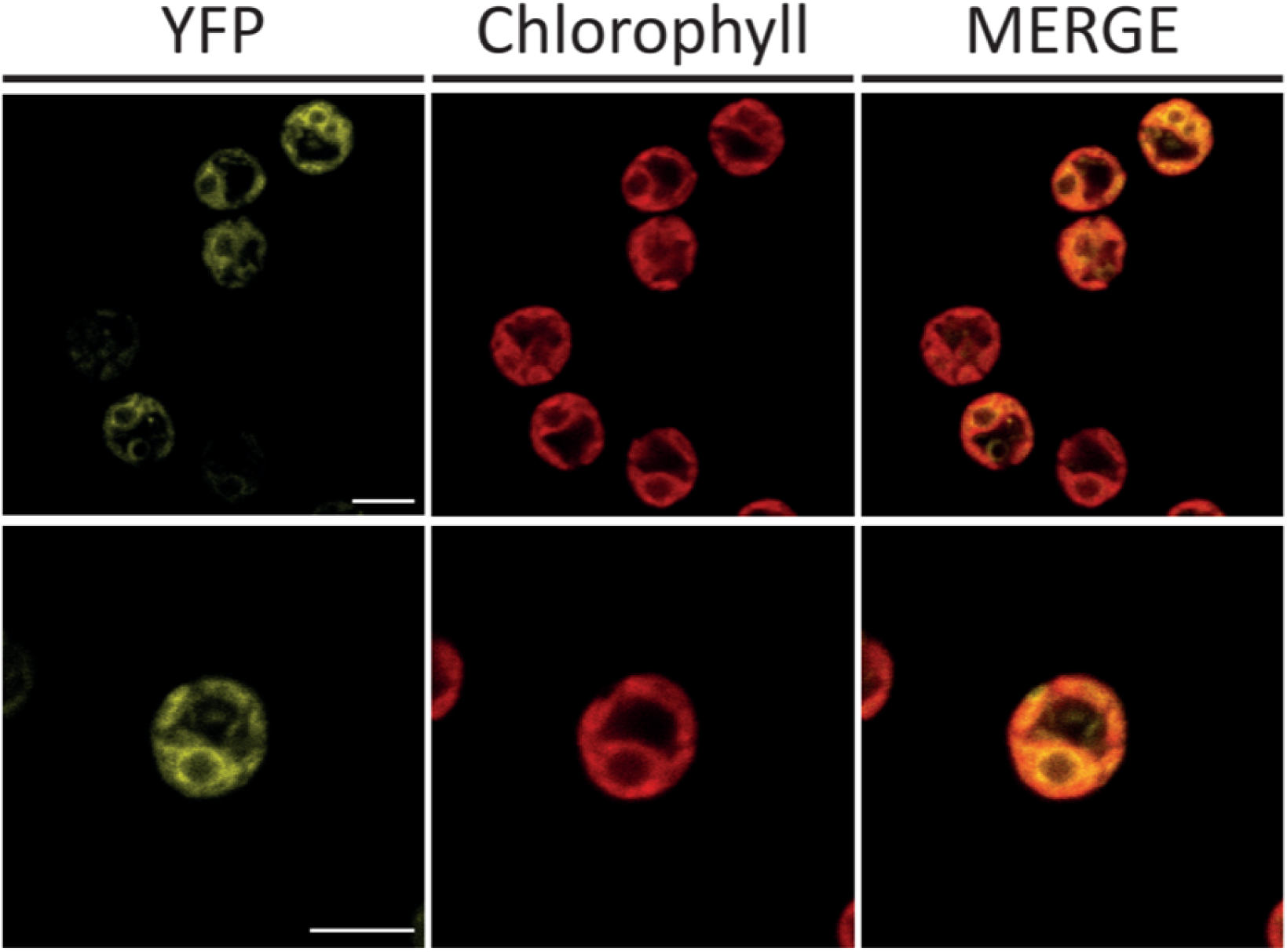
Analysis of FBP/SBPase localization. The presence of the fluorophore YFP fused to FBP/SBPase allowed for the investigation of the protein by confocal microscopy. YFP fluorescence (YFP), chlorophyll autofluorescence, and the merge of these two channels are shown. Scale bar represents 5 µm. Excitation 514 nm. Emission 522-572nm and 670-690 for YFP and chlorophylls, respectively.

### 3.2 Biochemical properties of dual-function FBP/SBPase in Chlamydomonas cells

To investigate the functional properties of the cyanobacterial FBP/SBPase, we purified the enzyme from *C. reinhardtii* cells by avidin-based affinity chromatography. By western blot analysis it was possible to confirm the purification of the recombinant protein with the expected molecular weight at 40 kDa (**Supplementary Figure 1**). The partially purified protein was then used to monitor the FBPase-related reaction in native conditions or following the reduction treatment (30 min) in the presence of 20 mM DTT (**Figure 3a and 3b**). The recombinant FBP/SBPase enzyme proved capable of catalyzing the dephosphorylation of FBP with a specific activity of 2.66 ± 0.73 µmol/min/mg, reaching a value of 4.75 ± 0.65 µmol/min/mg under reducing conditions (**Figure 3b and 3c**). The enzymatic activity was strictly dependent on magnesium ions, since no activity was monitored in the absence of MgSO_4_ (**Figure 3c**). By monitoring the activity response to variable substrate concentration, we extrapolated the Michaelis-Menten constant (*K*_m,FBP_), being 0.79 mM (**Supplementary Figure 2**). This result demonstrates that the FBP/SBPase enzyme expressed in *C. reinhardtii* can properly function as FBPase. Unfortunately, no *in vitro* assay is nowadays available to test the SBPase activity.

**Figure 3.**
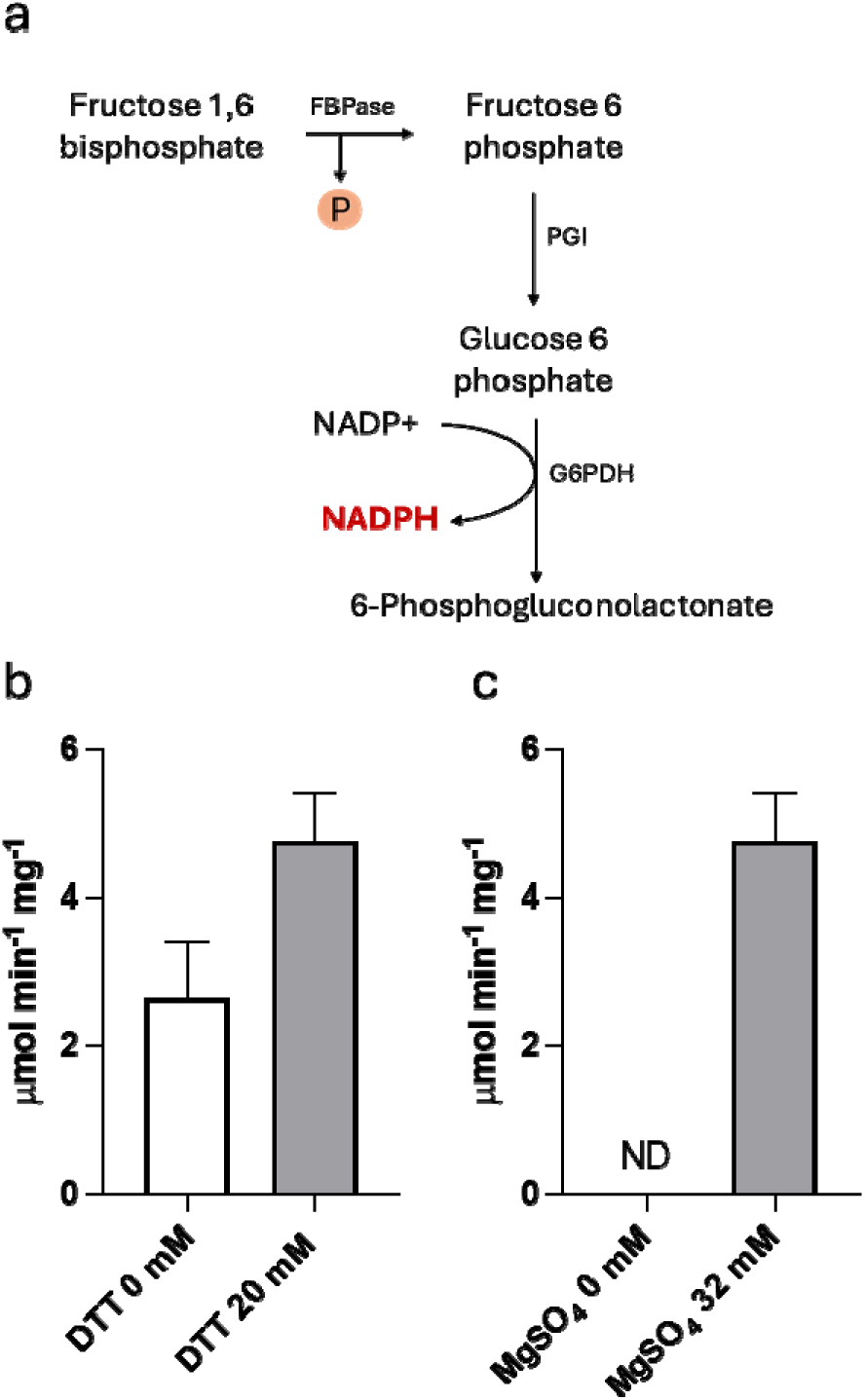
Kinetic analysis of cyanobacterial FBP/SBPase purified form Chlamydomonas cell. (a) Schematic representation of the enzymatic cascade used to assay FBPase activity. A coupled-enzyme assay including phosphoglucoisomerase (PGI) and glucose-6-phosphate dehydrogenase (G6PDH) was used to monitor FBPase activity by measuring the increase in absorbance due to NADPH production. (b) FBPase-related specific activity of the partially purified FBPase/SBPase in native (DTT 0 mM) or reducing (DTT 20 mM) conditions. (c) FBPase-related activity assayed with 0 or 32 mM of MgSO_4_ in the assay mix. Error bars are reported as standard deviations (n = 3).

### 3.3 Gene expression of endogenous FBPase and SBPase and heterologous FBP/SBPase genes in the transformant lines

The expression of dual-function FBP/SBPase and endogenous FBPases and SBPase were analyzed in the transformant lines and compared to the background. As reported in **Figure 4**, FBP/SBPase transcript was detected only in the transformant lines. In the case of endogenous SBPase, chloroplast FBPase and cytosolic FBPase no significant differences were measured in transcript content compared to the background strain upon normalization to the housekeeping gene Rack1 (**Figure 4**). The expression and activity of recombinant FBP/SBPase is thus not significantly influencing the gene expression of the endogenous FBPase and SBPase.

**Figure 4.**
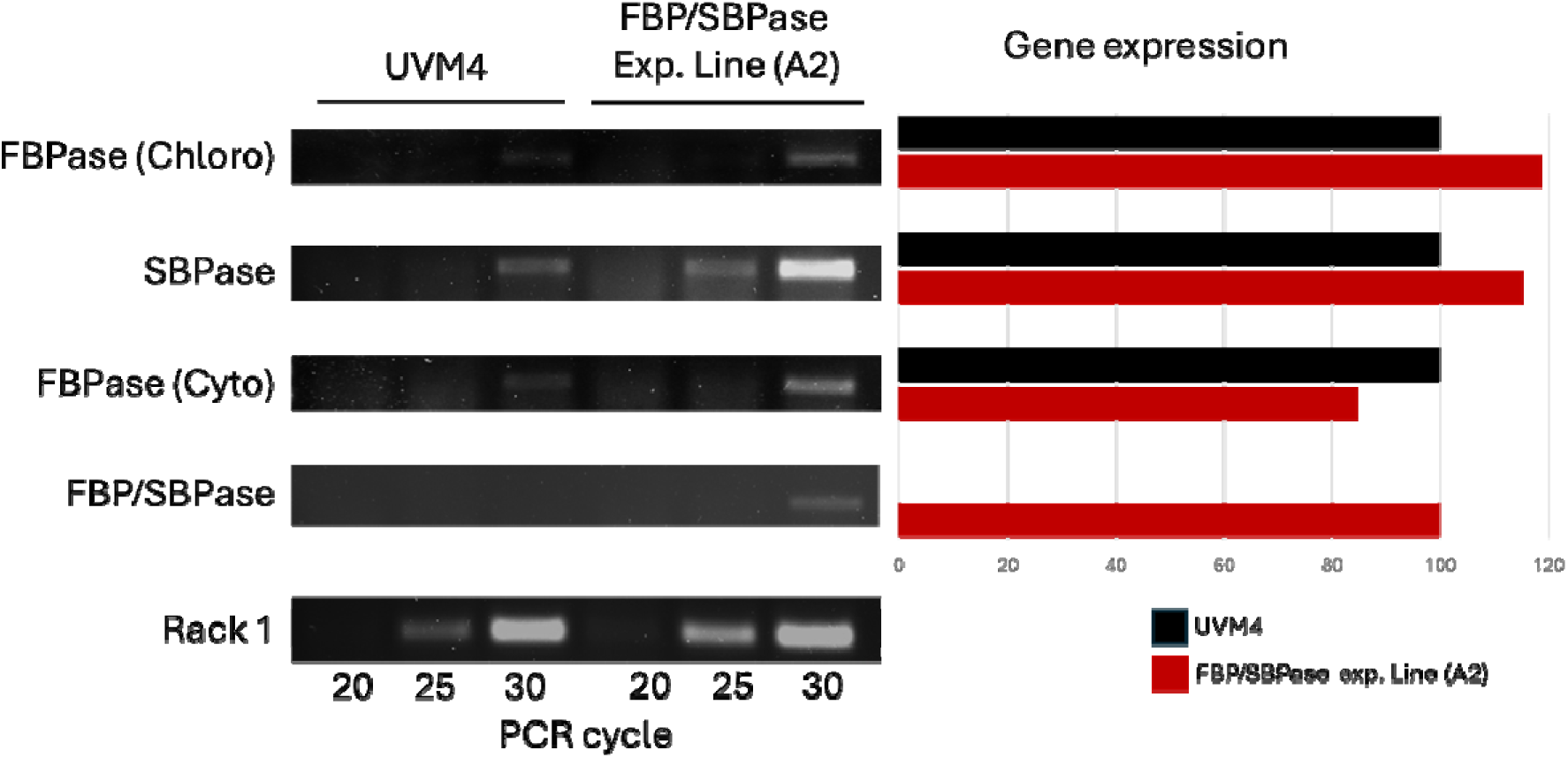
Gene expression of cyanobacterial FBP/SBPase and endogenous FBPases and SBPase. Gene expression of cytosolic FBPase, SBPase, chloroplast FBPase, FBP/SBPase, and the housekeeping rack1 was evaluated by semi quantitative RT-PCR. Relative gene expression with respect to UVM4 was estimated by densitometry on images of DNA agarose gel migration.

### 3.4 Consequences of heterologous FBP/SBPase expression on Chlamydomonas reinhardtii photosynthetic activity

The influence of cyanobacterial FBP/SBPase expression on photosynthetic activity of *C. reinhardtii* was then investigated in the transformant lines: photosynthetic rate can be determined by measuring the oxygen produced at different light intensities. As reported in **Figure 5**, increased net oxygen evolution was measured on a cell basis in FBP/SBPase expressing lines in cells grown either in photoautotrophic conditions (HS medium) or in mixotrophy (TAP medium), where acetate was adopted as the source of organic carbon. A significant increase in Pmax (being the maximum oxygen evolution rate) was measured in lines expressing FBP/SBPase compared to the background strain (**Supplementary Table 1**). This result suggests that on a cell basis, the catalytic activity of FBP/SBPase allows to increase electron transport from water to NADPH. However, it is worth noting that increased chlorophyll content per cell was measured in the FBP/SBPase expressing line compared to their background (**Supplementary Figure 3)**: when oxygen evolution data were normalized to chlorophyll content, no significant differences could be observed in the FBP/SBPase expressing lines (**Figure 5**).

**Figure 5.**
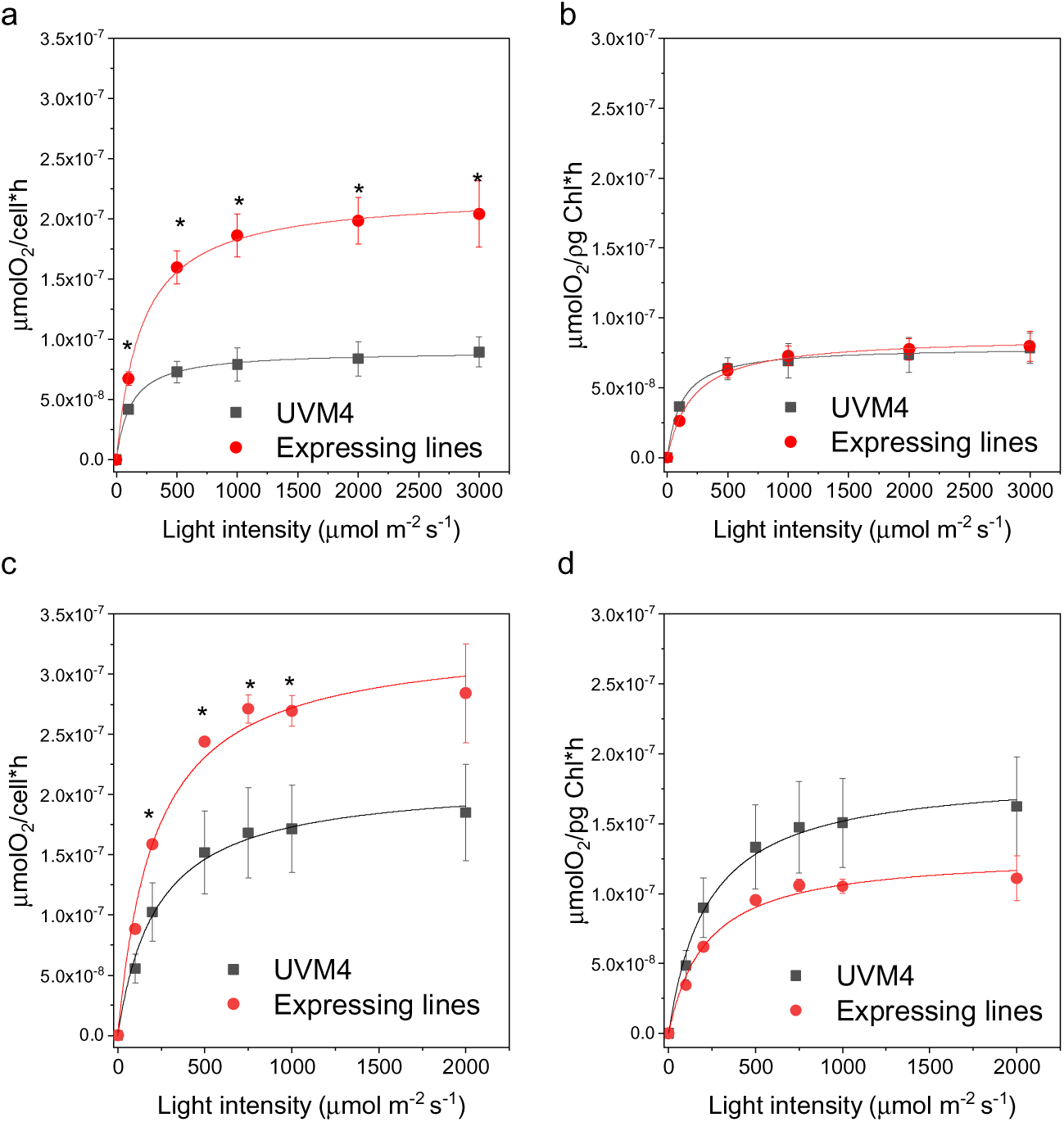
Light response and chlorophyl content of UVM4 and FBP/SBPase expressing lines. Light saturation curves of photosynthetic oxygen evolution obtained from UVM4 (black) and expressing lines (red) in mixotrophy (a,b) or photo autotrophy (c,d). Data are reported on cell basis (a,c) or on chlorophyll basis (b,d). Error bars are reported as standard deviations (n = 3). Asterisks indicate values that are significantly different compared to the UVM4 background strain (Student’s *t* test, *P* < 0.05).

Photosynthetic activity was then investigated measuring different photosynthetic parameters based on chlorophyll a fluorescence reflecting Photosystem II (PSII) activity. PSII operating quantum yield (Y(II)), electron transport rate (ETR), non-photochemical quenching (NPQ), and redox state of plastoquinone (1-qL) were analyzed in the transformant lines compared to the background strain. Y(II) is the effective PSII quantum yield at different actinic lights, while ETR is the relative charge separation rate at PSII reaction centers. NPQ describes the fraction of energy absorbed by PSII that is dissipated as heat by the activation of photoprotective dissipating mechanisms, while 1-qL indicates the fraction of closed PSII reaction centers, thus leading to plastoquinone reduction. As reported in **Supplementary Figure 4**, similar results were obtained for each photosynthetic parameter measured at different light intensities for the FBP/SBPase expressing line and the background strain. The results are consistent with similar oxygen evolution rates on a chlorophyll basis.

The accumulation of the main proteins involved in photosynthesis was then evaluated by western blot (**Figure 6**). CP43, PsaA, PetC, and AtpC subunits were analyzed to determine, respectively, PSII, PSI, cytochrome b6f, and chloroplast ATP synthase accumulation on a cell or chlorophyll basis. The antenna complexes of PSII, the LHCII proteins were also analyzed by using a primary antibody previously reported to recognize all the Lhcbm subunits found in the LHCII trimers ^67^. Finally, the content of RUBISCO and carbonic anhydrase 3 (CAH3) was investigated using specific antibodies (**Figure 6a**). As reported in **Figure 6b**, the densitometric analysis of the western blot results revealed that in the FBP/SBPase expressing lines a general increased accumulation on a cell basis for all the different protein subunits investigated, with statistically significant increase in the case of CP43, LHCII, PetC and RUBISCO compared to the background strain UVM4. Differentially, in the case of PsaA, CAH3, and AtpC, the increase on a cell basis in the transformant lines was not statistically significant. Considering the increased chlorophyll content per cell observed in the FBP/SBPase expressing lines, the results obtained were also plotted on a chlorophyll basis (**Figure 6c**). The CP43, LHCII and PetC content was similar on a chlorophyll basis in the FBP/SBPase expressing lines compared to the background strain, while PsaA, AtpC, RUBISCO and CAH3 content decreased. These results indicate that FBP/SBPase expression influences the content and organization of the photosynthetic apparatus, with relative increase of PSII and cytochrome b6f, while the ratio of RUBISCO/chlorophyll and RUBISCO/PSII decreased.

**Figure 6.**
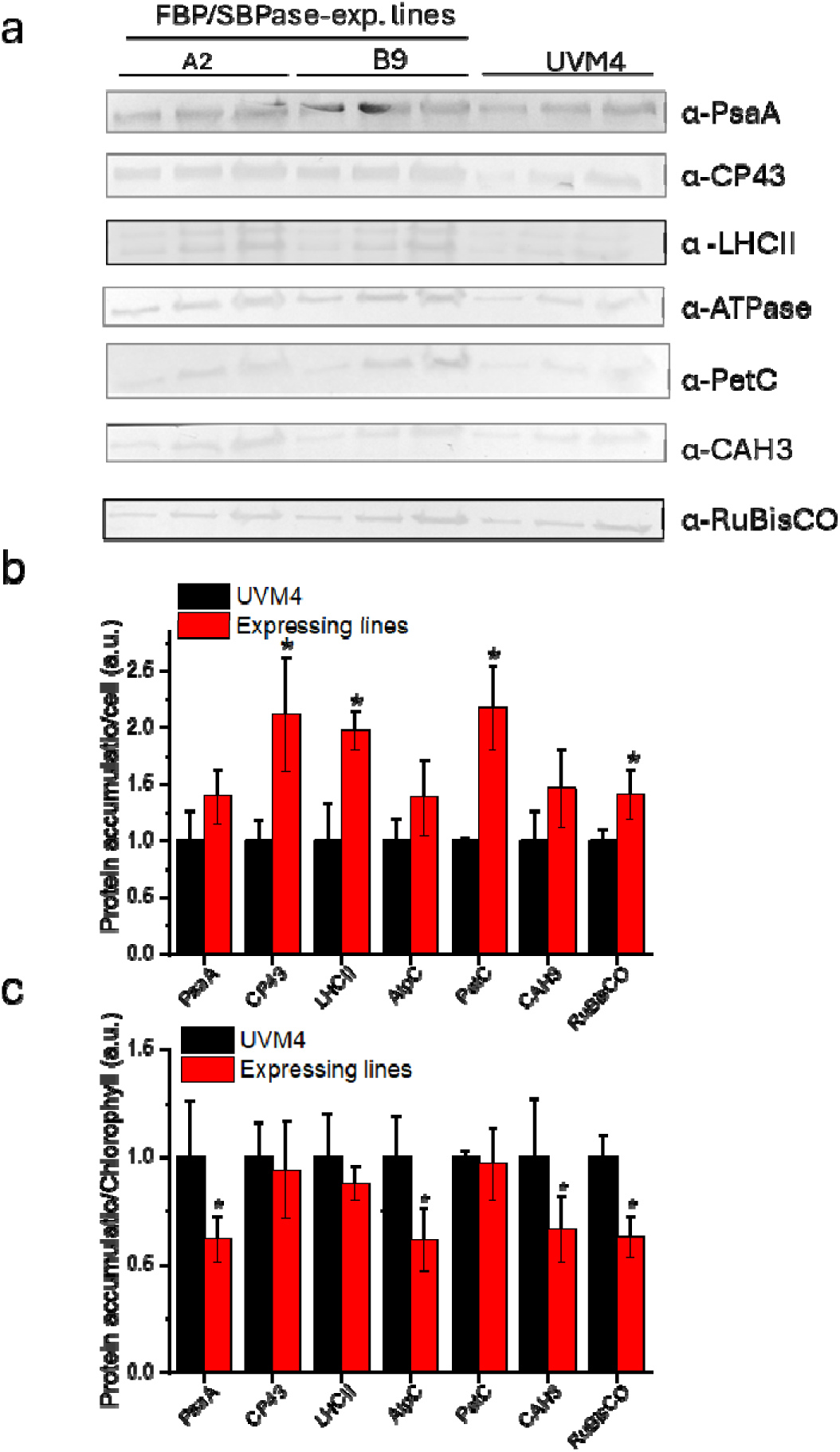
Immunoblot analysis of photosynthetic subunit accumulation. (a) Immunoblot analysis performed on UVM4 and FBP/SBPase expressing lines on total proteins extract using specific antibodies for PsaA, RuBisCO, ATPase C-subunit (AtpC), PetC, CP43, LHCII and carbonic anhydrase (CAH3). Different amounts of cells were loaded (3, 5 or 9 *10^6^ cells). Immunoblot signals reported in panel (a) were analyzed by densitometry to determine the relative protein abundance on cell basis (b) or on chlorophyll basis (c). Each protein level was normalized to the UVM4 protein level. Error bars are reported as standard deviations. Asterisks indicate values that are significantly different compared to the UVM4 background strain (Student’s t test, P < 0.05).

### 3.5 Dual-function FBP/SBPase expression improves resistance to photooxidation

The potential effect of improved CBB cycle activity on the photoprotective properties in the FBP/SBPase expressing lines was investigated by analyzing the kinetics of singlet oxygen production (**Figure 7**). Singlet oxygen is indeed produced by PSII upon exposure to strong light if the energy absorbed in excess is not safely dissipated or used by photochemical reactions ^68^. Single oxygen production can be analyzed upon exposure to strong red light using a fluorescent dye (Singlet Oxygen Sensor Green, SOSG) that increases its fluorescence after interaction with singlet oxygen, thereby enabling its quantification ^61^. Decreased SOSG fluorescence emission could be measured in the case of FBP/SBPase expressing lines compared to the background strain. This result is consistent with the increased Pmax on a cell basis upon expression of the cyanobacterial FBP/SBPase enzyme, suggesting an improved photosynthetic activity, and thus mitigating the risk of singlet oxygen formation. Interestingly, even on a chlorophyll basis, where similar Pmax were measured, decreased singlet oxygen formation was measured in the engineered lines. This is likely due to a faster replenishment of ADP and NADPL for the electron transport chain, which decreases the accumulation of harmful excited chlorophyll molecules.

**Figure 7.**
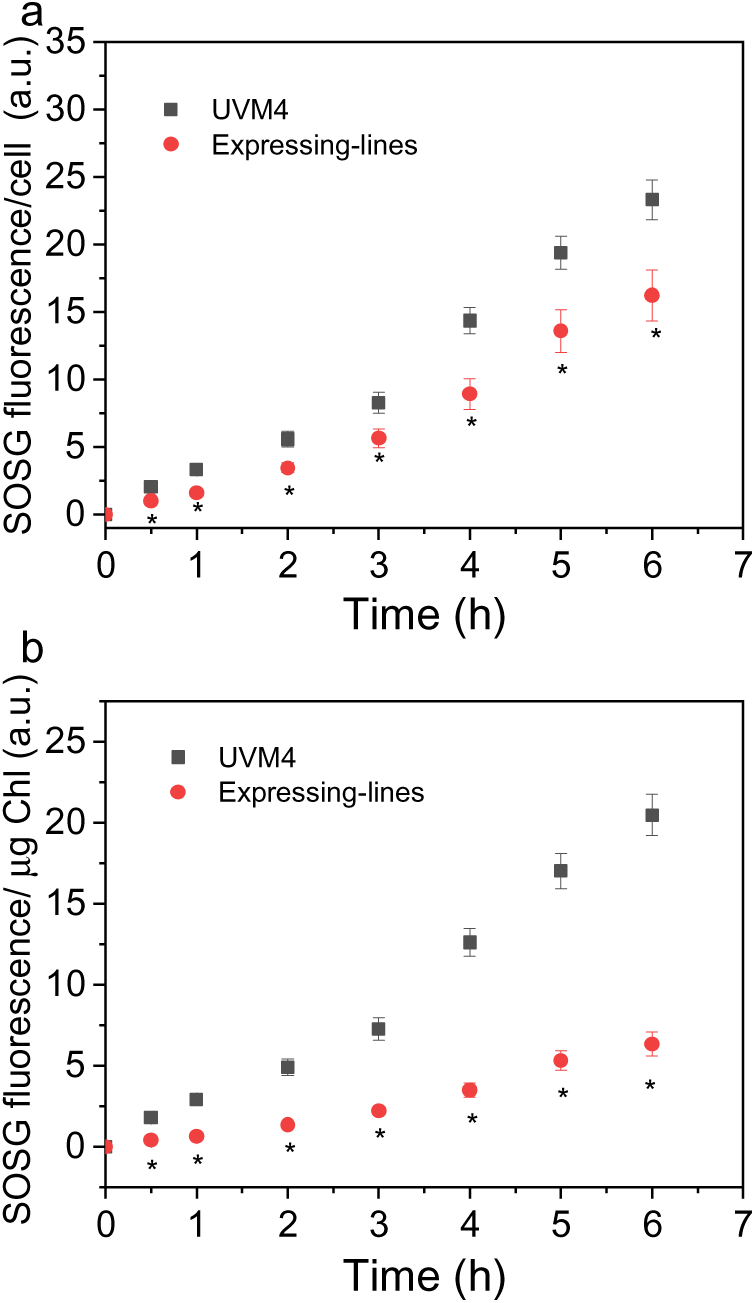
Singlet oxygen production for UVM4 and FBP/SBP expressing lines. Singlet oxygen production was estimated by fluorescence emission of Singlet Oxygen Sensor Green (SOSG) probe on cell basis (a) or on chlorophyll basis (b). Data reported are the mean values of three independent biological replicates with standard deviations indicated as error bars (n=3).

### 3.6 Consequences of heterologous FBP/SBPase expression on Chlamydomonas reinhardtii biomass production and composition

The possible influence of heterologous FBP/SBPase enzyme in *C. reinhardtii* was analyzed in small-scale airlifted photobioreactors. Two independent lines expressing FBP/SBPase and their UVM4 background strain were grown at 100 μmol photons m^−2^ s^−1^ in photoautotrophic or mixotrophic conditions. As reported in **Supplementary Figure 5**, similar growth curves and biomass accumulation were measured in cells grown in autotrophic conditions. Differently, engineered strains were characterized by a faster growth curve when cultivated in TAP medium (**Figure 8**). Despite the increased growth rate observed in the TAP medium, in both photoautotrophic and mixotrophic conditions, the engineered strains were characterized by a biomass concentration similar to that of the background at the end of the experiment. Cell density and cell size were then analyzed: a similar decrease in cell densities was observed in mixotrophic or photoautotrophic conditions (**Figure 8** and **Supplementary Figure 5**), whereas an increased cell volumes were measured in FBP/SBPase expressing lines compared to the background, either in photoautotrophic or mixotrophic conditions (**Figure 8**, **Supplementary Figure 5**, and **Supplementary Figure 6**).

**Figure 8.**
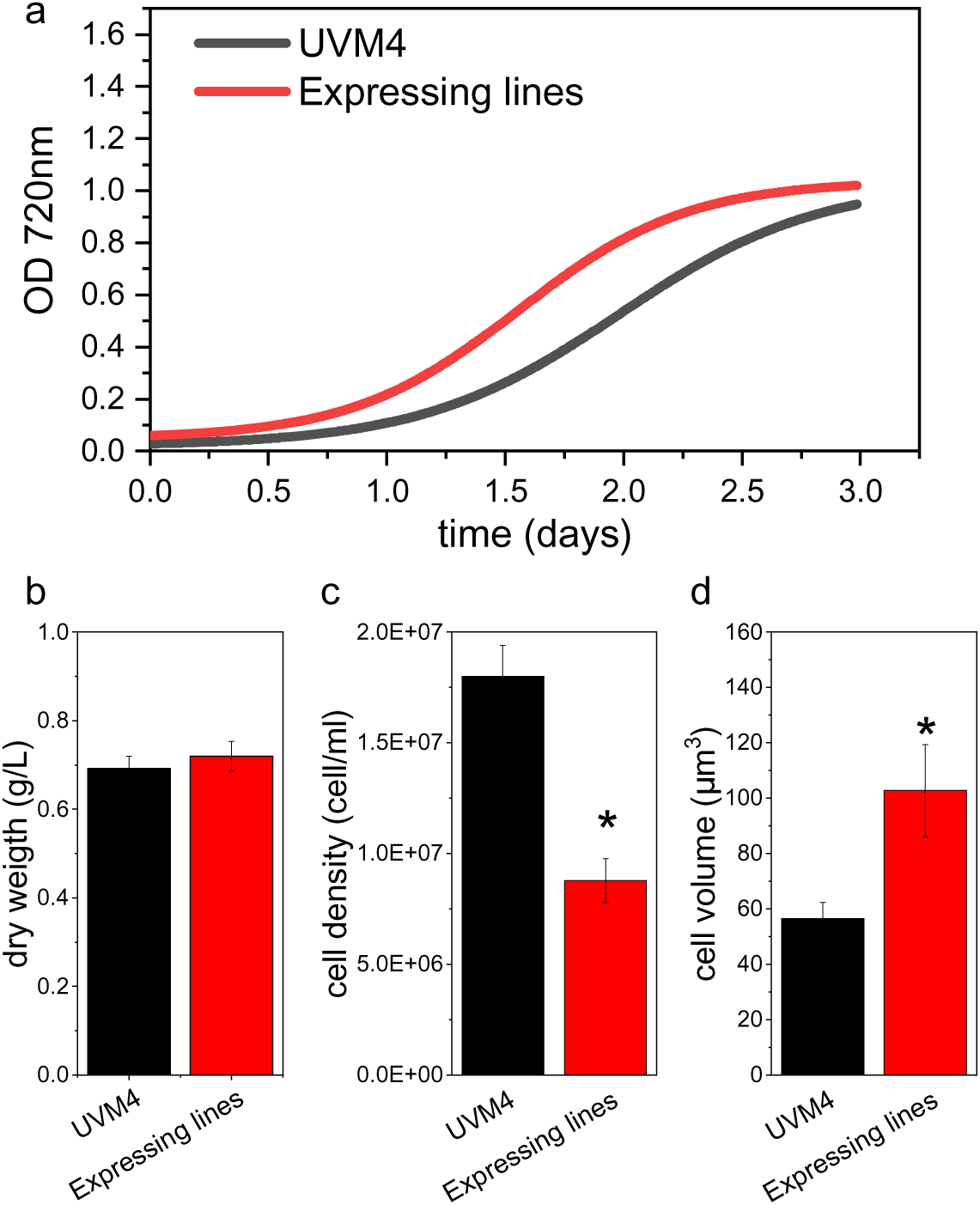
Biomass productivity of UVM4 and FBP/SBPase expressing lines. (a) Growth curves of UVM4 (black) and expressing lines (red) cultivated at 100 μmol photons m−2 s−1 in mixotrophy, monitoring OD at 720 nm. Volumetric biomass (b), cell density (c) and cellular volume (d). Error bars are reported as standard deviation (nL=L4). Data reported are the mean values of three independent biological replicates with standard deviations indicated as error bars. Asterisks indicate values that are significantly different compared to the background strains UVM4 (Student’s t test, P < 0.05).

Considering that most evident increase in growth curves were observed in TAP medium, additional conditions in mixotrophy were evaluated: transformant lines expressing FBP/SBPase and the UVM4 background strain were grown in atmospheric or 3% CO_2_-enriched air in low light (100 μmol photons m^−2^ s^−1^) or high light (1000 μmol photons m^−2^ s^−1^) conditions (**Supplementary Figure 7**). In low light conditions, only in the presence of low CO_2_ availability, increased growth kinetics were observed, while similar growth kinetics were measured upon exposure to high light, either in the presence of high or low CO_2_ availability (**Supplementary Figure 7**). As reported in **Supplementary Figure 8**, lower cell densities were measured in FBP/SBPase expressing lines grown at atmospheric CO_2_, exposed to either low or high irradiances. Cell volume was then measured for cells grown in different conditions, observing an increase in cell size in all conditions, not statistically significant only in the case of cells grown in low light in the presence of 3% CO_2_ (**Supplementary Figure 8**).

Altogether, these results suggest that the expression of the cyanobacterial FBP/SBPase affects cell dimensions when expressed in the chloroplast of *C. reinhardtii*, likely re-directing cell metabolism. At the end of the growth experiment, the biomass produced by cells grown in mixotrophy at 100 µmol photons m^−2^ s^−1^ was sampled to evaluate its composition, analyzing the main carbon sinks such as starch, proteins, and lipids. As shown in **Figure 9**, starch and protein content were increased in the FBP/SBPase expressing lines by a ∼2-fold factor compared to the background strain on a cell basis, while lipid content showed a 50% increase on the same basis. This is consistent with the increased cell volume observed in the transformant lines. Considering the fraction of starch and protein on dry weight, a 20-25% increase was observed for both compounds in the FBP/SBPase expressing lines, even if this increase was not statistically significant. While a 50% increase on cell basis was observed in teh FBP/SBPase expressing lines also in the case of lipids, no major differences was observed for the lipid fraction of cell dry weight.

**Figure 9.**
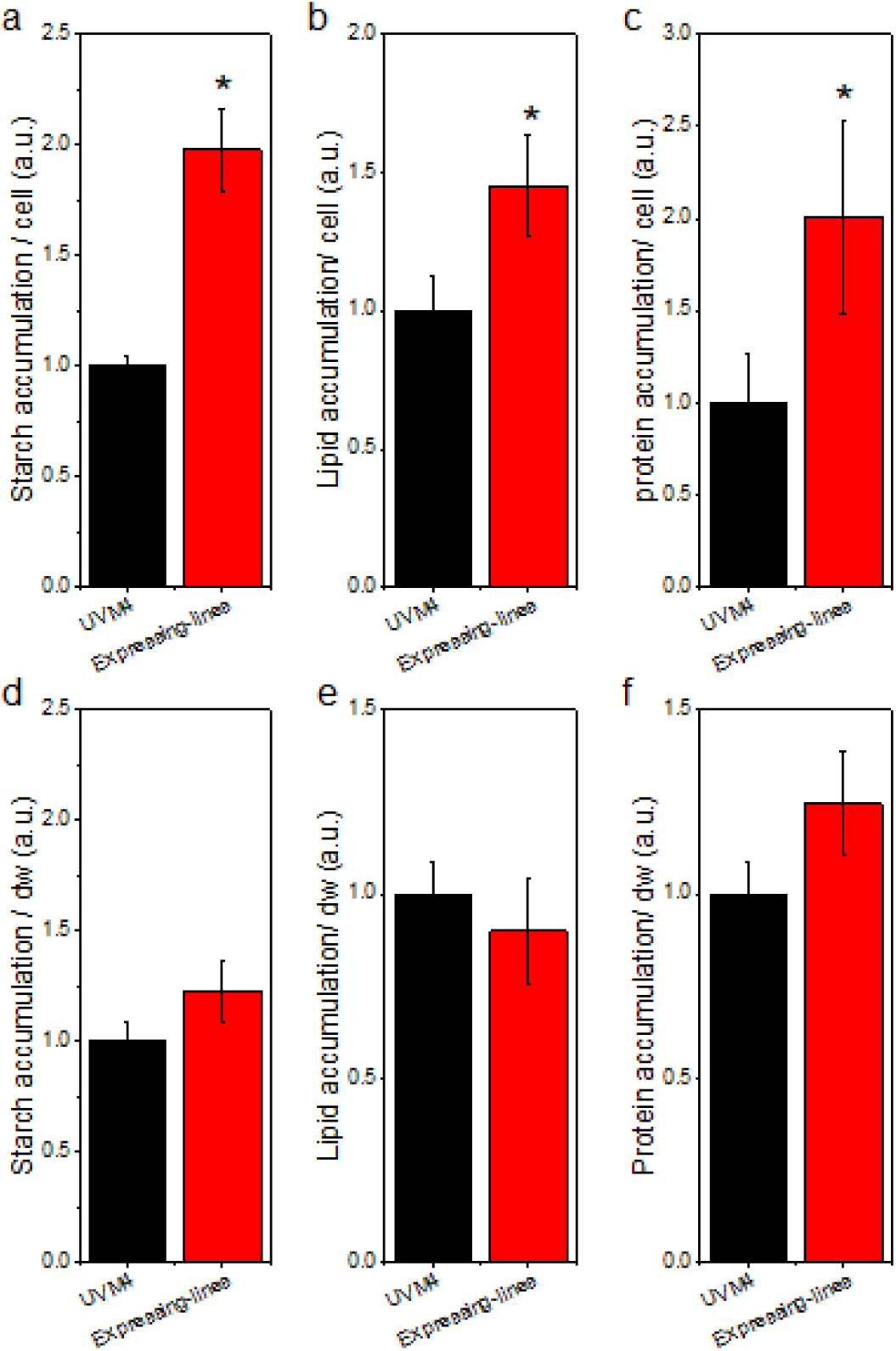
Effect of FBP/SBPase expression on starch, lipid and protein accumulation. Starch (a,d), polar lipid (b,e) and protein accumulation (c,f) on cell basis (a,b,c) or on dry weight basis (d,e,f). Error bars are reported as standard deviations (n=4). Asterisks indicate values that are significantly different compared to the background strain UVM4 (Student’s t test, P < 0.05).

Interestingly, the increased cell volume measured in the case of FBP/SBP expressing lines caused an increased sedimentation rate (**Figure 10**). For spherical cells, sedimentation is described by Stokes’ law and depends on the cell’s size and the density of both the cells and the liquid ^69^. According to this model, the increased volume of the FBP/SBPase expressing cells is estimated to cause a ∼60% higher sedimentation rate, which is the result of ∼80% higher buoyant mass. This is consistent with the experimental data (**Figure 10**), where the measured sedimentation rate in expressing lines was 57% higher compared to the UVM4 case.

**Figure 10.**
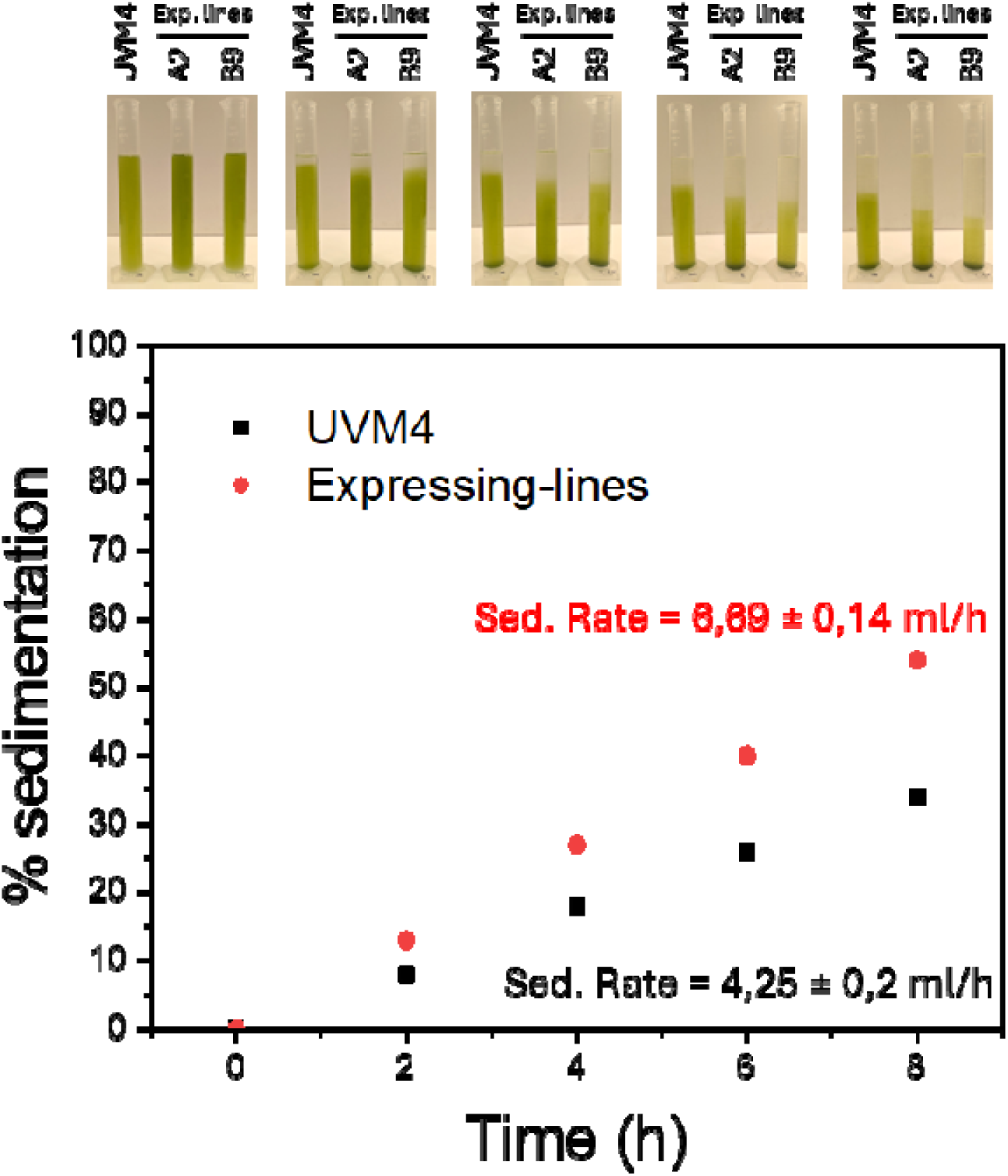
The sedimentation percentage for UVM4 (black) and expressing lines (red) was determined by measuring the volume of clarified liquid over time in a 50 mL graduated cylinder. The sedimentation rate, expressed as the volume of clarified liquid (mL) per hour, is shown in black for UVM4 and in red for the expressing lines.

This phenotype of the FBP/SBPase expressing line could lead to easier and less costly harvesting processes of the biomass produced.

## 4. DISCUSSION

Cyanobacterial FBP/SBPase can simultaneously catalyze two key steps of the CBB cycle (*i.e.*, FBPase- and SBPase-mediated reactions), which are involved in regeneration phase ^41^. These reactions were previously identified as limiting steps for CBB cycle and have been previously targeted for biotechnological manipulation to improve carbon fixation ^12,24,25,40,70^. A more efficient CBB cycle leads to increased consumption of ATP and NADPH and more rapid regeneration of ADP, phosphate and NADP^+^, which are needed to desaturate the light phase of photosynthesis. Recombinant expression of the dual-function FBP/SBPase enzyme in the chloroplasts of transgenic plants of *Nicotiana tabacum* leads to improved photosynthesis accompanied by increased carbohydrate accumulation and biomass production ^28^. Similar results were obtained by heterologous expression of this FBP/SBPase enzyme in lettuce, soybean, rice, and in the microalga *Euglena gracilis* ^42–45^. Overexpression of these two CBB enzymes in the model green alga *C. reinhardtii* had complex phenotypes, leading in some cases even to detrimental effects, as in the case of FBPase overexpression ^38^. On the contrary, the overexpression of endogenous SBPase resulted in an increased photosynthetic rate under conditions of high light and elevated CO_2_ concentrations ^10^. Based on these previous results, the expression of cyanobacterial FBP/SBPase enzyme was tested in *C. reinhardtii*. This microalga has also been recognized as safe by the FDA in the USA and its dried biomass powder is intended to be used as a nutritive ingredient in food to replace other dietary proteins ^71^. Enhancing the carbon fixation efficiency of *C. reinhardtii* could increase its photosynthetic rate and, consequently, the biomass yield, which can be utilized across a wide range of industrial applications.

The FBP/SBPase enzyme was expressed in *C. reinhardtii*, adopting the strategy of synthetic gene redesign, taking advantage of the enhanced transcription led by the insertion of RUBISCO SMALL SUBUNIT 1 intron ^47^. Recombinant FBP/SBPase enzyme was correctly localized in the chloroplast as demonstrated by confocal microscopy on lines obtained by transformation with the PsaD_FBP/SBPase_YFP construct (**Figure 2**). Considering the possible negative effect of YFP fusion on the catalytic activity of the FBP/SBPase protein, the FBP/SBPase-YFP fusion protein was only considered for the localization experiments. The recombinant FBP/SBPase protein was isolated from the engineered lines and proved to be correctly folded and active by *in vitro* activity test. It is worth noting that only FBPase activity can be monitored *in vitro*, as the substrate for SBPase enzyme is currently unavailable. Even the FBP/SBPase dual enzyme required the presence of Mg^2+^ for its catalytic activity, which was also regulated by its redox state, being full reduction of the protein necessary for full activity of the protein *in vitro* (**Figure 3**).

The expression of FBP/SBPase in the chloroplast caused an improved photosynthetic activity on a cell basis. Indeed, light-dependent oxygen evolution rate increased consistently with the increased content of chlorophyll and components of the light phase of photosynthesis in the engineered strains (**Figure 5**). The oxygen evolution rate was not different on a chlorophyll basis, in line with the similar ETR, Y(II), 1-qL, and

NPQ measured in the engineered cells compared to the background **(Figure 5** and **Supplementary Figure 4).** The expression of cyanobacterial FBP/SBPase thus allowed the cell to increase the light phase of photosynthesis, accumulating more photosynthetic pigments and more chlorophyll-binding proteins, causing an increased capacity to transport electrons from water to NADPH and consequently produce ATP. ATP and NADPH constitute key molecules for the proper functioning of the CBB cycle (or other metabolic pathways) and their recycling is critical to prevent saturation of the photosynthetic apparatus ^12^. The increased PLLL observed on a per-cell basis in the FBP/SBPase-expressing lines indicates that ATP and NADPH cofactors are effectively utilized by the CBB cycle, thereby preventing saturation of the photosynthetic light reactions. The expression of FBP/SBPase caused a decreased accumulation of singlet oxygen, the main ROS species produced at the level of PSII when exposed to saturating light ^68^, suggesting an increase tolerance to photooxidative stress likely thanks to improved CBB cycle and improved regeneration of NADP^+^, ADP, and free phosphates. Interestingly, the RUBISCO/chlorophyll and RUBISCO/PSII ratios were decreased upon expression of FBP/SBPase: it was previously demonstrated that the RUBISCO/PSII ratio is a limiting factor for the photosynthetic activity in cyanobacteria ^19^. The amount of RUBISCO affects the maximum capacity for carboxylation/oxygenation, depending on the substrate availability. It was indeed previously reported that high CO_2_ availability decreases the RUBISCO content, while the opposite occurs in CO_2_ limitation ^60^. The decreased RUBISCO content on a chlorophyll basis could thus be related to increased availability of one of its substrates, whose accumulation depends on the regeneration phase of the CBB cycle.

The consequence of FBP/SBPase expression was an increased growth rate in mixotrophic conditions when both CO_2_ and light are limiting factors. The similar growth rate and biomass accumulation in photoautotrophy even upon expression of FBP/SBPase indicates that the CBB is not sufficiently boosted in the engineered lines to increase biomass productivity. Nevertheless, in both mixotrophic and autotrophic growth, an increased cell size was observed upon expression of the FBP/SBPase enzyme. The increased cell size in FBP/SBP expressing lines was specifically accompanied by increased protein, lipid, and starch content on a cell basis, with starch and protein also slightly increased as a fraction of dry weight. Increased starch content is likely related to increased sugar content in the chloroplasts of the transformant lines, thereby enabling starch accumulation. The availability of reduced organic carbon in mixotrophic conditions decreases the requirement of triose-phosphate export from the chloroplast to the cytosol, and in the FBP/SBPase expressing lines, this led to an increased availability of CBB cycle products in the chloroplast to be used for starch synthesis. The increased protein content can be related again to improved CBB cycle in the FBP/SBPase expressing lines, which also boost the cytoplasmic carbon metabolism, where the acetate availability can be redirected toward amino acid biosynthesis rather than being used for oxidative catabolism. In general, the boosted CBB cycle caused by FBP/SBPase activity results into a significant increase in biomass when it is merged with the acetate-based metabolism allowing to use a greater fraction of organic carbon from acetate and/or from photosynthates to support mainly starch and protein accumulation. High CO_2_ availability and/or high light was indeed overcoming the positive effect obtained by overexpression of FBP/SBPase. Either CO_2_ availability and irradiance were indeed reported to affect the CBB cycle activation: high CO_2_ availability and/or exposure to strong light could thus trigger the endogenous CBB cycle enzymes, making ineffective the overexpression of the cyanobacterial FBP/SBPase. Finally, the presence of a regulation point cannot be excluded: in vitro activity test demonstrated that the cyanobacterial FBP/SBPase required a reducing agent to be fully active (**Figure 3**). The redox control of cyanobacterial FBP/SBPase could potentially limit its activity in the engineered cells, decreasing the consequent effect on biomass productivity.

It is worth noting that the increased cell volume of the FBP/SBPase expressing lines leads to more prominent cell sedimentation **(Figure 10**). This phenotype is relevant for industrial cultivation of *C. reinhardtii* and microalgae in general, being the harvesting processes one of the most energy and cost-demanding steps in the overall cultivation process ^72^. The expression of FBP/SBPase enzyme could thus be considered as a strategy to improve the cell harvesting process, allowing the inclusion of a preliminary step of cell sedimentation, which is currently not effective in the wild-type strains of *C. reinhardtii*.

In conclusion, the expression of the FBP/SBPase enzyme in *C. reinhardtii* allowed for the improvement of the CBB cycle, leading to enhanced biomass, starch, and protein productivity in mixotrophic conditions, when CO_2_ and light are limiting. CBB cycle is finely regulated with several enzymes and molecules that come into play potentially representing additional limiting steps that remain to be investigated. For this reason, further studies in plants, microalgae, and other transgenic organisms are needed to better understand the regulatory mechanisms that interact in photosynthesis and that may lead to the development of transformed lines with improved photosynthetic activity and increased biomass or metabolite yield of interest.

## DATA AVAILABILITY

All data generated or analyzed during this study are included in this published article and its supplementary information files.

## ACKNOWLEDGEMENTS

We thank the Centro Piattaforme Tecnologiche for providing access to the core facilities of the University of Verona. The research was supported by the ERC Starting Grant SOLENALGAE (679814) to M.B. and by the Italian Ministry of University and Research (MUR) grant PRIN 2022 (2022ASSR9R) to M.B. and M.Z.

## AUTHOR CONTRIBUTIONS

**Martina Bussola:** Investigation, Data curation, Validation, Visualization, Writing - review & editing; **Federico Perozeni:** Investigation, Data curation, Validation, Methodology, Supervision, Writing - review & editing; **Maria Meloni, Matteo Pivato:** Investigation, Data curation, Methodology, Writing - review & editing; **Mirko Zaffagnini:** Conceptualization, Funding acquisition, Supervision, Methodology, Validation, Visualization, Writing - review & editing; **Matteo Ballottari:** Conceptualization, Funding acquisition, Supervision, Methodology, Validation, Visualization, Project administration, Writing - original draft, Writing - review & editing.

