## Supplementary Figure for "Heterologous expression of the cyanobacterial fructose-1,6-/sedoheptulose-1,7-bisphosphatase in *Chlamydomonas reinhardtii* causes increased cell size and biomass productivity in mixotrophic conditions"

**Supplementary Figure 1. FBP/SBPase purification from *Chlamydomonas reinhardtii* expressing lines.** Immunoblot against Strep-Tag II (A) and Coomassie blue stained gel (B) performed on the FBP/SBPase purified using the Strep-Tactin XT resin column. (C) Quantitative SDS-PAGE for quantification of the recombinant FBPase/SBPase using CrFBPase as a standard.

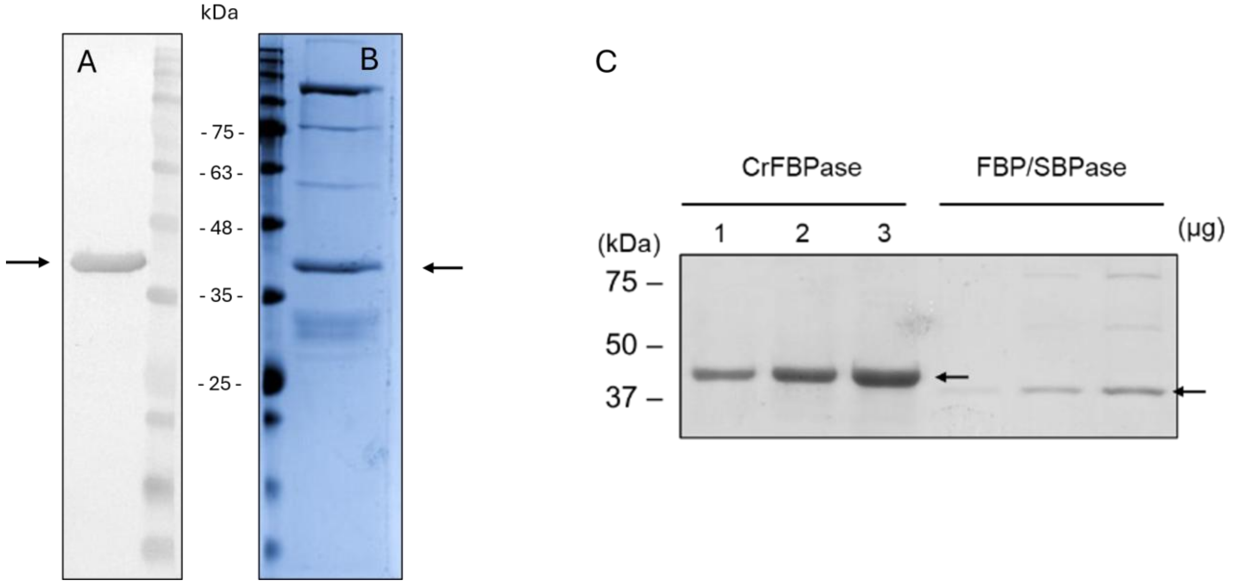

**Supplementary Figure 2.** ***In vitro*** **analysis of FBPase catalytic activity of recombinant FBP/SBPase**. A representative experiment of dose-dependent effect of fructose-1,-6-phosphate (FBP) on the activity of purified FBP/SBPase from expressing lines is reported.

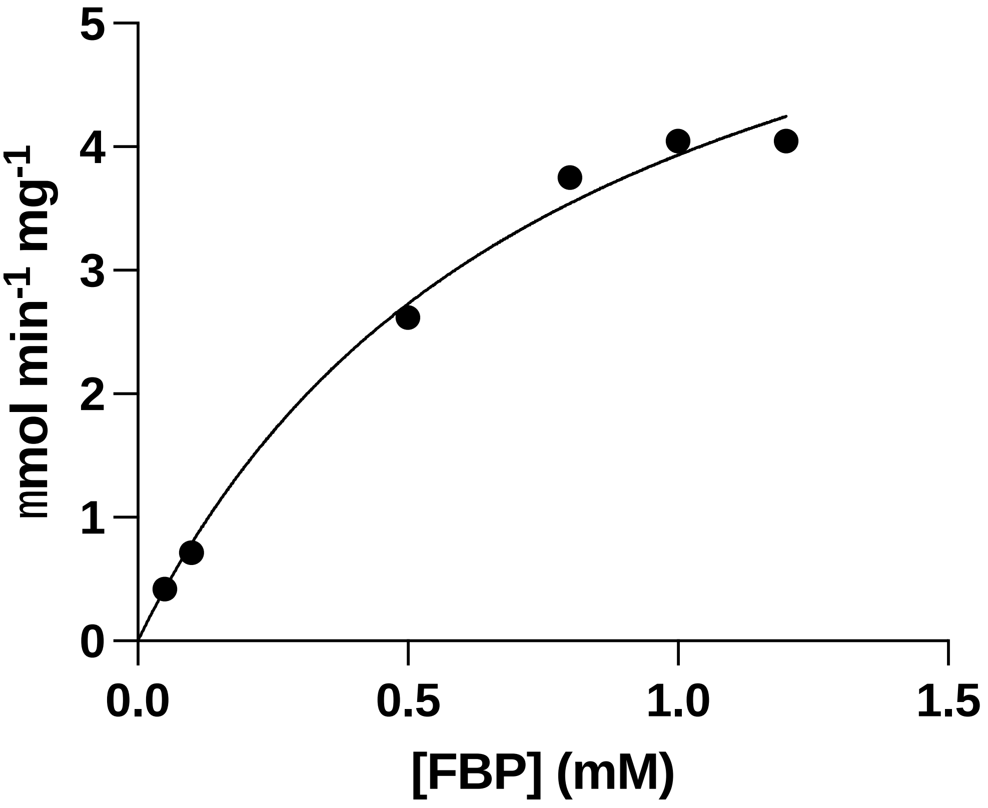

**Supplementary Figure 3. Chlorophyll content per cell in FBP/SBPase expressing lines compared to the background case**. Chlorophyll content on cell basis in UVM4 (black) and expressing lines (red) (b).

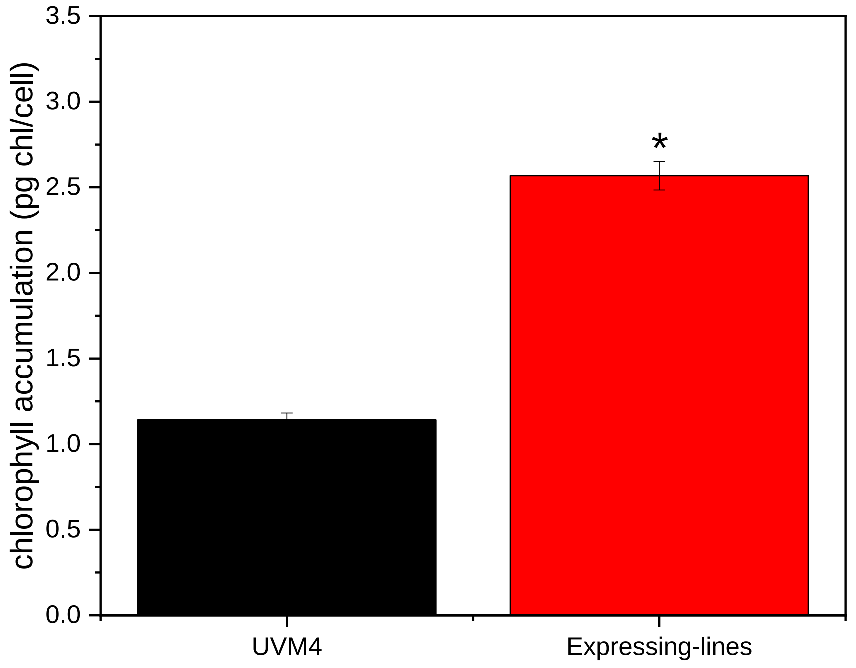

**Supplementary Figure 4. Impact of FBP/SBPase expression on Photosystem II photochemical and non-photochemical activity**. (A) PSII operating quantum yield (Y(II)), (B) non-photochemical quenching (NPQ) measured at different actinic lights in dark-adapted cells, (C) PSII electron transport rate (ETR(II), (D) redox state of plastoquinone (1-qL). Data reported are means of three biological replicates with standard deviation shown.

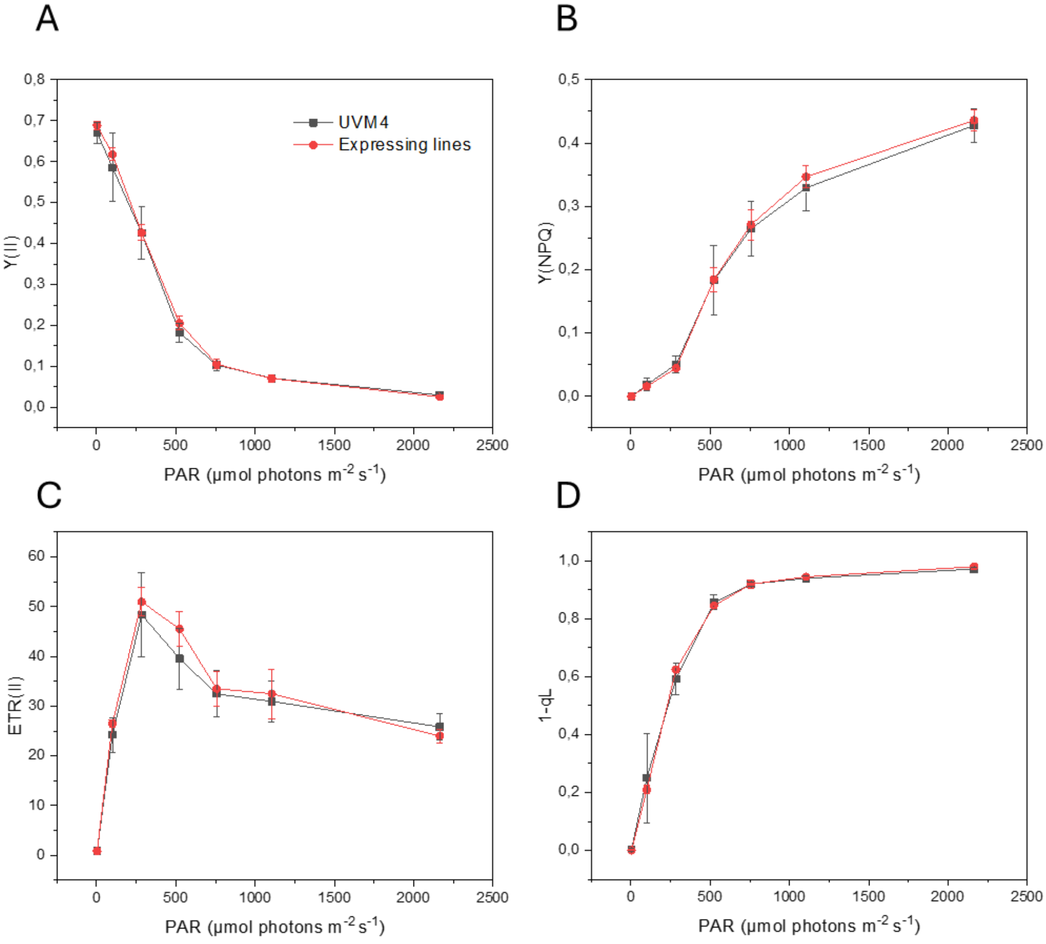

**Supplementary Figure 5. Biomass productivity of UVM4 and FBP/SBPase expressing lines.** Growth curves of UVM4 (black) and expressing lines (red) cultivated at 100 µmol photons $m^{-2}$ $s^{-1}$ in photoautotrophy, monitoring OD at 720 nm (A). Volumetric biomass (B), cell density (C) and cellular diameter (D). Error bars are reported as standard deviations (n = 4). Values significantly different compared to UVM4 are reported with * (Student's t test, $P < 0.05$)

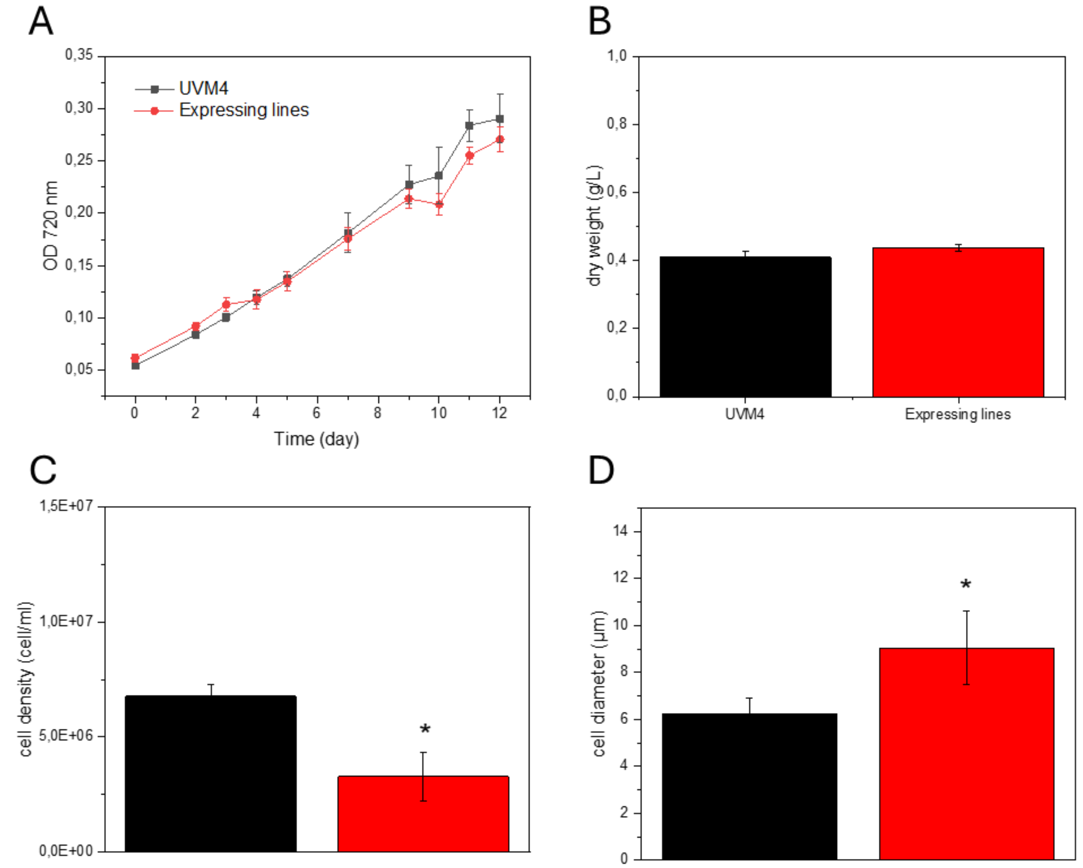

**Supplementary Figure 6. Morphology of FBP/SBPase expressing cells.** Microscopy images of background and engineered cells. Images of UVM4 (left) and FBP/SBPase expressing lines (A2 and B9, right) grown in TAP medium at exponential phase are shown with scale bar (5 μm).

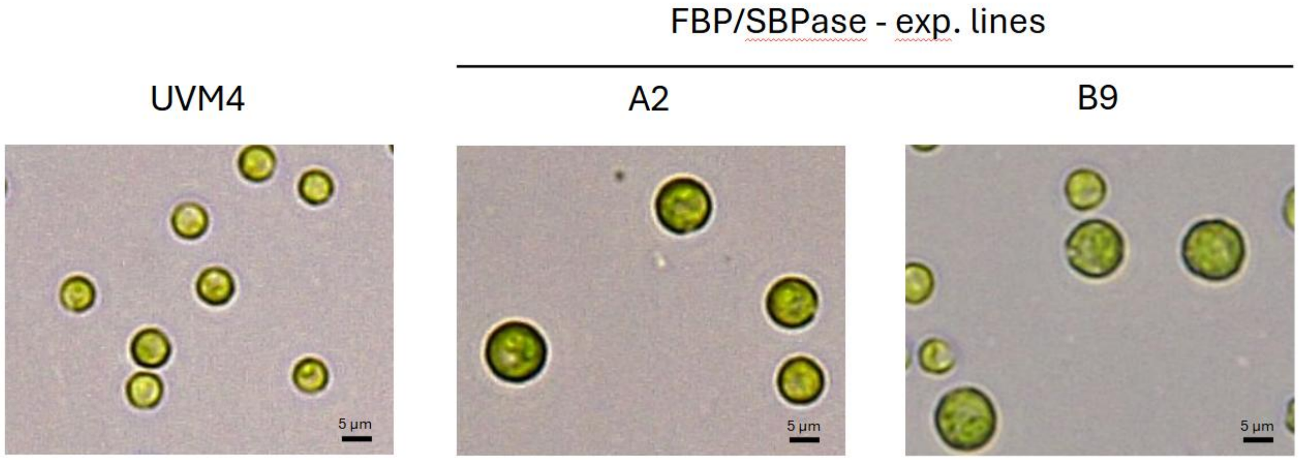

**Supplementary Figure 7**: **Effect of FBP/SBPase expression on algal growth in mixotrophy.** Growth curve generated by following cell scattering at 720 nm and volumetric productivity for UVM4 (black) and FBP/SBPase expressing lines (red). Two different light intensities were used: 100 µmol photons m$^{-2}$ s$^{-1}$ for low light (LL) and 1000 µmol photons m$^{-2}$ s$^{-1}$ for high light (HL) with air with atmospheric or 3% enriched $CO_2$ concentration. Error bars are reported as standard deviations (n = 4).

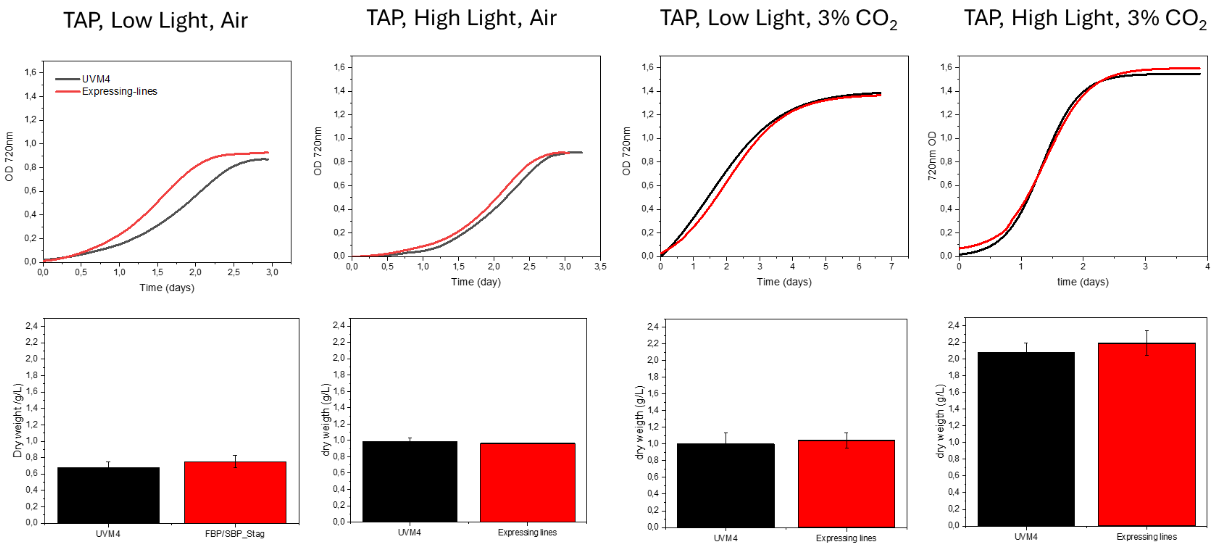

**Supplementary Figure 8. Effect of FBP/SBPase expression on cell density and dimension**. Cell density (A) and cellular volume (B) of UVM4 (black) and FBP/SBPase expressing lines (red) at the end of growth reported in Supplementary Figure 7. Error bars are reported as standard deviations (n = 4). Values significantly different compared to UVM4 are reported with * (Student's t test, $P < 0.05$)

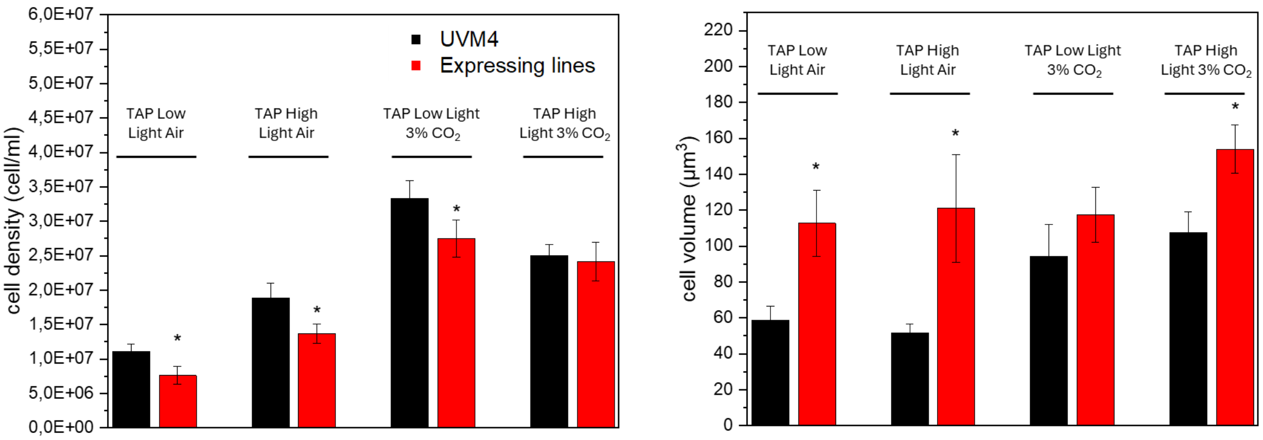


**Supplementary Table 1. Fitting results of oxygen evolution curves.** Net oxygen evolution data were obtained upon subtraction of oxygen consumption rate in the dark and fitted with the hyperbolic function y Pmax * x/(KI + x), where Pmax is the maximum net oxygen evolution rate and KI the light intensity at which the net oxygen evolution rate is Pmax/2.

| | | UVM4 | | EXPRESSING LINES | |
|---|---|---|---|---|---|
| TAP | Pmax (µmolO2/cell*h) | 9.01E-08 | 1.27E-09 | 2.21E-07* | 3.36E-09 |
| | Pmax (µmolO2/rg Chl*h) | 7.90E-08 | 1.12E-09 | 8.63E-08 | 1.31E-09 |
| | KI (µmol $m^{-2}$ $s^{-1}$) | 118.4 | 9.9 | 208.7* | 15.7 |
| HS | Pmax (µmolO2/cell*h) | 2.12E-07 | 7.04E-09 | 3.31E-07* | 1.46E-08 |
| | Pmax (µmolO2/rg Chl*h) | 1.95E-07 | 9.47E-09 | 1.51E-07 | 8.82E-09 |
| | KI (µmol $m^{-2}$ $s^{-1}$) | 225.3 | 27.2 | 215.6 | 35.3 |
